# Dynamics of EEG Microstates Change Across the Spectrum of Disorders of Consciousness

**DOI:** 10.1101/2024.05.30.596582

**Authors:** Dragana Manasova, Yonatan Sanz Perl, Nicolas Marcelo Bruno, Melanie Valente, Benjamin Rohaut, Enzo Tagliazucchi, Lionel Naccache, Federico Raimondo, Jacobo D. Sitt

**Affiliations:** Sorbonne Université, Institut du Cerveau - Paris Brain Institute - ICM, Inserm, CNRS, Paris 75013, France; Université de Paris Cité, Paris, France; Department of Information and Communication Technologies, Centre for Brain and Cognition, Computational Neuroscience Group, Universitat Pompeu Fabra, Barcelona, Spain; Department of Physics (University of Buenos Aires), Buenos Aires, Argentina; National Scientific and Technical Research Council (CONICET), Buenos; - AP-HP, Hôpital de la Pitié Salpêtrière, Neuro ICU, DMU Neurosciences, Paris, France; AP-HP, Hôpital Pitié-Salpêtrière, Service de Neurophysiologie Clinique, Paris, France; Institute of Neuroscience and Medicine (INM-7: Brain and Behaviour), Research Centre Jülich, Germany; Institute of Systems Neuroscience, Heinrich Heine University Düsseldorf, Germany

**Keywords:** Disorders of Consciousness, EEG microstates, Dynamics, Entropy production

## Abstract

As a response to the environment and internal signals, brain networks reorganize on a sub-second scale. To capture this reorganization in patients with disorders of consciousness and understand their residual brain activity, we investigated the dynamics of electroencephalography (EEG) microstates. We analyze EEG microstate markers to quantify the periods of semi-stable topographies and the large-scale cortical networks they may reflect. To achieve this, EEG samples are clustered into four groups and then fit back into each time sample. We then obtain a time series of maps with different frequencies of occurrence and duration. One such occurrence of a map with a given duration is called a microstate. The goal of this work is to study the dynamics of these topographical patterns across patients with disorders of consciousness. Using the microstate time series, we calculate static and dynamic markers. In contrast to the static, the dynamic metrics depend on the specific temporal sequences of the maps. The static measure Ratio of Total Time covered (RTT) shows differences between healthy controls and patients, however, no differences were observed between the groups of patients. In contrast, some dynamic markers capture inter-patient group differences. The dynamic markers we investigated are Mean Microstate Durations (MMD), Microstate Duration Variances (MDV), Microstate Transition Matrices (MTM), and Entropy Production (EP). The MMD and MDV decrease with the state of consciousness, whereas the MTM non-diagonal transitions and EP increase. In other words, DoC patients have slower and closer to equilibrium (time-reversible) brain dynamics. In conclusion, static and dynamic EEG microstate metrics differ across consciousness levels, with the latter capturing the subtitler differences between groups of patients with disorders of consciousness.

**Abbreviated summary:** This study investigates EEG microstate dynamics in patients with disorders of consciousness (DoC) to understand residual brain activity and network reorganization. Entropy production, in addition to static markers, differs between patients and healthy controls, whereas the rest of the dynamic markers show differences across patient groups.

## Introduction

Consciousness is hypothesized to arise out of complex network interactions on a sub-second scale ^1,2^. Various theoretical models build on the idea that conscious states do not rely on a single cortical area or network but require brain-wide communication^1^. In patients with Disorders of Consciousness, the network coordination is altered ^2^, and thus the characterization of these pathological network dynamics can aid diagnostic and prognostic analyses. Addressing this challenge, Demertzi and colleagues^3^, have demonstrated that different recurrent patterns of brain activity, obtained from resting-state fMRI, are directly associated with disorders of consciousness (DoC) clinical categories. Particularly, one of the phase-coherence patterns more present in higher global states of consciousness is characterized by long-range functional communication between brain areas^3^.

The exploration of resting-state brain activity relies on the theory that the resting-state networks reflect an inner state of exploration which optimizes the system for input and thus it influences perception and cognitive processing ^4^. However, to respond to changes in the environment, networks must reorganize on a sub-second time scale. Unlike fMRI, the EEG temporal resolution can capture fast fluctuations. Using high-density EEG while tracing the ongoing brain activity of patients with a disorder of consciousness (DoC), we employ a method known as EEG microstates as a proxy to track latent brain state changes on a sub-second scale.

EEG microstates are defined as successive short periods (around 50-100 ms) during which the configuration of the scalp potential field remains semi-stable^5,6^. As reported in the literature^6–11^, four prototypic maps can be reliably identified across healthy participants. These four maps are commonly denoted with letters and have distinct topographical descriptions: map A has a left-right orientation, map B has a right-left orientation, C has an anterior-posterior orientation, and D has a frontocentral maximum. The temporal characteristics of microstates are proposed to represent the basic building blocks of spontaneous mental processes, as well that the quality of mentation is determined by their occurrence and temporal dynamics^6^. Furthermore, EEG microstates have been investigated in neurological and clinical conditions such as narcolepsy, Alzheimer’s, disorders of consciousness, and schizophrenia^6,9,12^. The dominance of certain maps in time has also been associated with the activation of an underlying resting state network^13^.

To assess the dynamics of the microstates in patients with a DoC compared to healthy controls, we used the EEG recordings of the auditory local-global paradigm^14^. The DoC clinical diagnostic categories with which we work in this paper are the Unresponsive Wakefulness Syndrome (UWS) (also known as the Vegetative State - VS), the Minimally Conscious State (MCS), and the Emergent Minimally Conscious State (EMCS)^15–17^. Recent work has also investigated the use of EEG microstates in the classification of DoC patients^12,18,19^. They show that some microstate markers can aid the DoC classification, as well as the patients’ outcome prediction. However, neither study investigates the dynamic microstate markers in detail.

The first working hypothesis of this study is focused on the investigation of the microstate dynamics across DoC classes and aims to test whether similar to the loss of consciousness in sleep there is a lengthening of the microstates across DoC patients. In other words, UWS patients will have significantly longer microstate durations than MCS, EMCS, and HC. Furthermore, we hypothesized that the variance of microstate durations would be highest for HC and lowest for UWS. Another approach to assess the change of dynamics is the concept of time-reversibility^20^, in which we assess whether the discrete state transitions are symmetrical when going from brain state A to brain state B and reversed. Recent work has shown that entropy production can be assessed from resting-state and task data and can be used to differentiate between global states of consciousness^20,21^. The microstates discretization of the whole-brain activity, allows us to test a measure of entropy production. We hypothesized that the transition probabilities matrices of the healthy controls would show a lack of symmetry and that the transition entropy would scale from low to high for patients in UWS to the healthy controls.

## Methods

### Participants

The study involved two groups of participants: a healthy group used as a control for the analyses (n=37, mean age 27.6 +/− 5.8, 8 female), and a group of patients with disorders of consciousness. The healthy participants took part in the study voluntarily as approved by the ethical committees at the Paris Brain Institute (ICM), Salpetriere, Sorbonne University (Inserm CPP C13-632 41) and by the Comité de Protection des Personnes Ile-de-France. The patients’ recordings were acquired as part of an operational diagnostic procedure at the Neurology Department at the Salpetriere Hospital or as part of other studies aiming for the development of diagnostic and prognostic analyses^22,23^. According to the behavioral assessment by a neurologist using the CRS-R, the patients were classified to be in Unconscious Wakefulness Syndrome (UWS) otherwise known as Vegetative State (VS) (n=70, mean age 46.4 +/− 18.1, 22 female), Minimally Conscious State (MCS) (n=70, mean age 43.7 +/− 18.6, 28 female), Emergent MCS (EMCS) (n=14, mean age 38.9 +/− 23, 4 female).

### Experimental design and data characteristics

The EEG data were recorded with a 256-electrode geodesic sensor net (EGI®, Oregon, USA) referenced to the vertex. The sampling rate was set to 250 Hz. For the goal of this study, we used recordings during a Local-Global (LG) Paradigm^14^. Specifically, we used the period of the paradigm during which four identical sounds are presented. We refer to this period as a pseudo-resting state as it elicits a non-specific response^22^.

### Data pre-processing

Python 3.7-based open-source software libraries such as the MNE-Python^24^ and the NICE-tools and extensions^22^, were used for the EEG signal processing. The recordings were downsampled to 250 Hz, band-pass filtered (0.5 - 45 Hz) then segmented in epochs ranging from −200 ms to +1344 ms relative to the first sound onset. The data was epoched to enable better artifact cleaning where single strongly artefacted epochs were removed. Similar approaches have been taken in other studies^25,26^. Electrodes with voltages exceeding 100 µV in more than 50% of the epochs were removed. Moreover, voltage variance was computed across all correct electrodes. Electrodes with a voltage variance Z-score higher than 4 were also removed. This process was repeated four times. Bad electrodes were interpolated using a spline method. Epochs were labeled as bad and discarded when the voltage exceeded 100 µV in more than 10% of electrodes. The remaining stimulus-locked epochs were re-referenced to an average reference. In addition, the channels along the edges of the head were removed systematically from all the recordings. This was done due to the electrodes containing face muscle artifacts or not having sufficiently strong signals. In the cases where more of the 50% of the epochs were marked as bad, the subject was then taken out of the analysis. They are not included in the patient summaries above.

### Microstates Analysis

EEG microstates have been studied for more than 30 years^5,6^. Nevertheless, there are still ongoing debates on the subtleties regarding their derivation and analysis. In this study, a combination of the procedures most widely accepted in the scientific community was followed^6–8,10–12,27–29^. We used a modified version of the MNE microstates sub-package as well as metrics introduced initially in other studies^10,11^.

#### Microstates clustering and segmentation methods

The Global Field Power (GFP) of the signal is calculated which quantifies the variance of voltage potentials across all of the electrodes^30^. As a measure of the strength of the scalp potential at a given time point, the GFP is based on the potential differences between all electrodes. The output is a vector of scalar values per sample. In the past, high GFP has been associated with stable EEG topographies^7,27^. The maps were traditionally seen as discrete, meaning they do not gradually morph into one another or overlap in time, but rather a single map is dominant and then abruptly transitions to another map^5^. However, this has been disputed in recent studies^31,32^. To obtain the time series of microstates, the EEG epochs from a single participant were stacked going from a 3D array (epochs, channels, samples) to a 2D array (channels, samples). Next, all the data samples are z-scored by taking into account the mean and the standard deviation of the whole recording for each channel. For our goal, we extracted the EEG potential fields at the GFP maxima (the minimal peak distance was set to 2 samples), also referred to as maps (Figure 1). The topographies were filtered according to two criteria. The first filtering was done to remove the maps at GFP peaks that were larger than one standard deviation above the GFP mean (as suggested in ^28^). Secondly, the maps that belong to a GFP local maxima but within the lowest 15% of the GFP signal were removed. As shown by Mishra et al.^31^, in this GFP range, the k-means clustered maps are equidistant from the EEG potential fields at the given sample. The remaining GFP peaks were subsampled randomly to less than 100.000 maps per recording. This was done to optimize the computations. The remaining maps were submitted to a modified k-means clustering algorithm as described by Pascual-Marqui et al.^30^. The modified k-means clustering algorithm and the segmentation functions were available as a sub-package of MNE Python^24^. The specificity of this algorithm is that it is polarity-invariant, meaning that topographies with opposite polarity are assigned to the same class^30^. Not all clustering algorithms are polarity invariant, one such example is classical k-means clustering. In this thesis, the number of clusters (k) was set to four to compare the results to existing literature, as in most studies this is the chosen number of clusters^6–9,12^.

**Figure 1.**
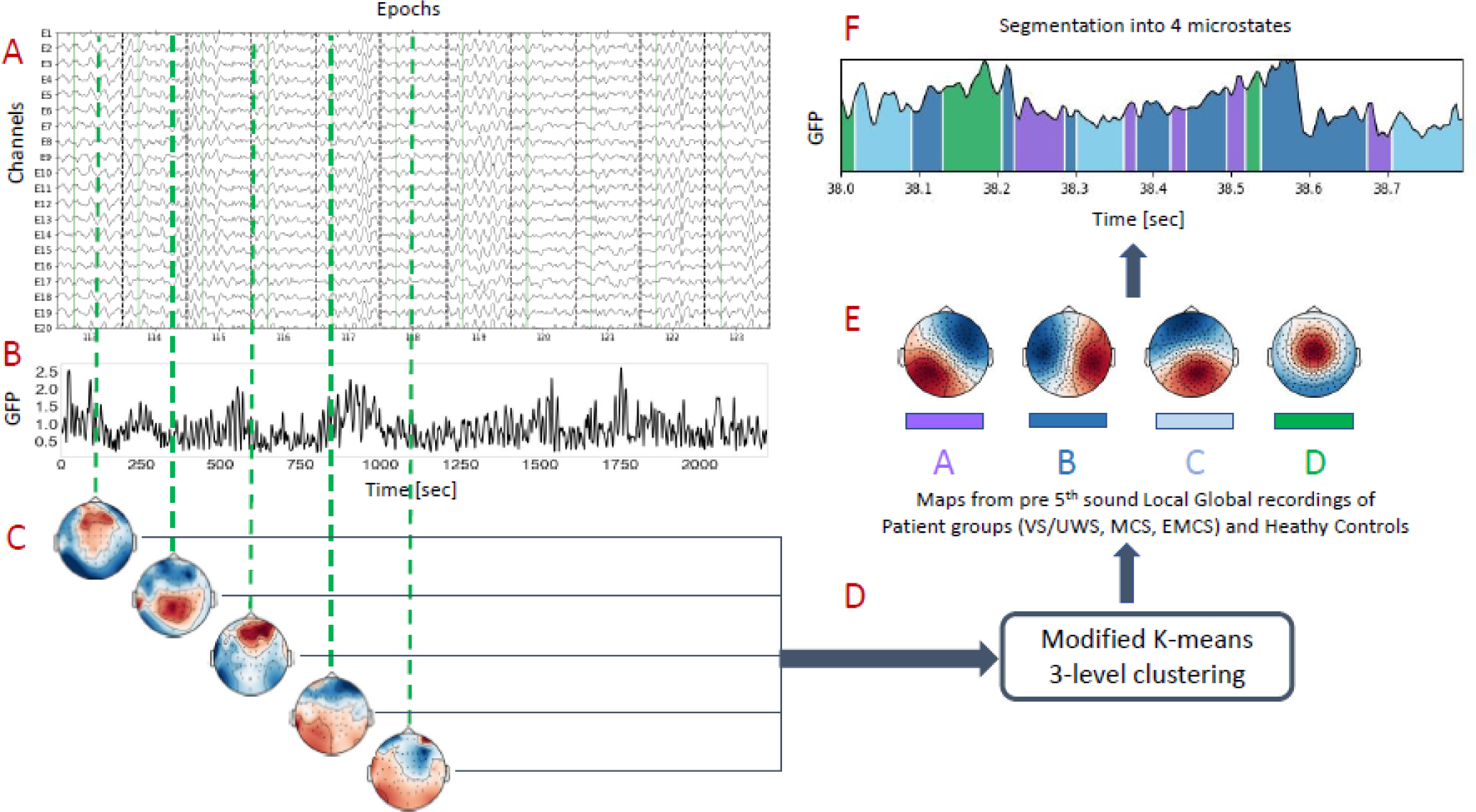
Illustration of the microstates clustering and segmentation algorithms. **(A)** The participants’ EEG is divided into epochs (a sample of epochs and channels is shown). **(B)** The Global Field Potential (GFP) is calculated by taking the EEG signals from all the electrodes at each time point. **(C)** The GFP peaks are extracted and filtered according to two criteria (explained in the Materials and Methods). The topographies (EEG potentials from all the electrodes) are extracted at each GFP peak. **(D)** The extracted topographies at the GFP peaks are used as input to a modified k-means clustering algorithm. **(E)** The output of the clustering gives a k-number of topographies that explain most of the variance in the data. Each topography or microstate map is denoted with a different color - shown in the square below the maps. **(F)** The maps are fitted back to the EEG channel signal to obtain the sequence of dominant maps along with the duration of the recording, this is known as segmentation. Abbreviations: Global Field Power (GFP), seconds (sec), Vegetative State (VS), Unresponsive Wakefulness Syndrome (UWS), Minimally Conscious State (MCS), Emergent Minimally Conscious State (EMCS).

To assess to what extent the maps explain the data, a measure called Global Explained Variance (GEV) is used (Equation 1). The first step is to fit the map closest to the EEG field potential at the given time point. This is done for each sample of the EEG signal and the output 1D vector (map per sample) is denoted as a segmentation. In other words, the maps are back-fitted to new EEG samples based on topographical similarity. The segmentation gives the information on which map is closest to the topography sample by sample. The topographical similarity is calculated based on a distance measure called Global Map Dissimilarity (GMD)^33^. This measure looks at how similar the topography maps are and is invariant to the strength of the signal. In other words, two maps with similar topographies, but different EEG voltage strengths will result in a low GMD distance^33^.

GEV is a measure of the similarity of each EEG sample to the microstate map it has been assigned to. It is calculated by the multiplication of the squared correlation between the EEG sample and the assigned map with the sample’s fraction of the total squared GFP^33^. The k-means clustering is randomly initiated 100 times, and at each iteration, the GEV and the corresponding maps are saved. This is done to maximize the GEV by selecting the maps from the iteration with the highest GEV value.

For a group-level analysis, we do a three-level clustering (Supplementary Figure 3). First, we clustered into 10 maps on a subject level. Next, because of the unbalanced dataset (eg. 70 MCS subjects versus 14 EMCS), we perform bootstrapping. In other words, we sample with repetition the groups. The sample size is equal to the smallest group size, in our case it is 14 as we have only 14 subjects with EMCS. In each bootstrap iteration, we cluster the subject-level maps into four maps. We repeat this 2000 times. We obtain an array of 4 x 2000 maps, which are clustered into 4 maps. The final four maps are the ones we use to fit back to the subject time series to obtain the per-subject segmentation. When a given map is dominant over a few continuous (uninterrupted) samples, this is what we call a microstate. Using this time series (the segmentation) we calculate the microstates markers. A similar but simpler double clustering procedure has been previously used^26,29^.

Another method used to deal with EEG noise is to smoothen the segmentation. Due to various short-lived artifacts of a few samples, the back-fitting of the maps to the EEG can be affected. For this reason, the segmentation is smoothened using a window smoothing algorithm explained in Titterington et al.^34^ (introduced in Pascual-Marqui et al.^30^ and implemented in the MatLab EEG microstates toolbox by Poulsen et al.^28^).

#### Microstates markers

For the goal of this study, we investigated markers calculated from the microstate segmentation^6,7,10^. We divided the microstate markers into static and dynamic. The static metrics are time-independent, meaning they are not influenced by the specific temporal sequence of the microstates. They are:

- Ratio of Total Time covered (RTT), sometimes referred to as segment count density, or empirical symbol distribution^10,11^. It denotes the fraction of the total time for which a certain microstate is dominant. In other words, it indicates the percentage of time covered by a given microstate map over the duration of all the epochs.
- Global Explained Variance (GEV) per map reflects the ratio of variance explained by each of the k-group maps - calculated for all the epochs of one participant. This reflects the goodness of fit of the microstates to describe the map sequence of that subject. The markers GEV and RTT depend on one another. Meaning that the bigger the RTT coverage is, the more variance the given map is likely to explain within the data. Contrastly, the GEVs of the maps reflect the topographical similarity to each data point (how well they correlate), whereas the RTT accounts for the discrete temporal presence without reflecting the correlation between the dominant map and the given sample^9^.

The dynamic markers which depend on the temporal sequence of map alterations are:

- Mean Microstate Duration (MMD) which represents the mean of the microstate durations in milliseconds per participant. This marker gives information on the stability of the microstates and can allow for the comparison of whether microstate transitions occur faster or slower in a given group of participants.
- Microstate Duration Variance (MDV) represents the variance of the microstate durations per participant. It reflects the microstates’ duration variability, and their temporal consistency or lack thereof. In other words, it shows whether the durations of the microstates within a subject are consistently long, or consistently short, or if they have a variable length.
- Microstates Transition Matrix (MTM) describes the transition between the k-microstates. It denotes the ratio of the number of transitions from one microstate map to another and is calculated from the transitions present in all the epochs from a single participant’s EEG recording. The transition probabilities quantify how frequently a given map is followed by the other maps. It characterizes the flow of microstates in terms of particular sequences of transitions and allows one to determine if this sequence has a particular dynamical order.
- Entropy Production (EP) quantifies the symmetry of the transition probabilities. In other words, it is a measure of broken detailed balance (where the transition probability of going from map i to map j Pij is different than the transition probability going from map j to map i Pji and this difference leads to EP increase). This analysis has been implemented in previous fMRI studies in resting state and task paradigms^20,21^.

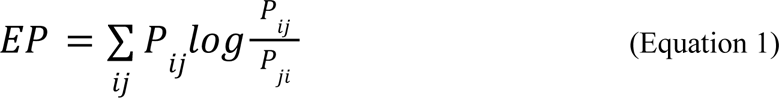

### Statistical analyses

Due to the microstate marker distributions per group being skewed, we performed non-parametric statistical tests. To investigate the differences per pair of groups, the Mann-Whitney U test was used for comparing independent data samples. This test is a nonparametric version of the independent samples t-test. Its null hypothesis stipulates that the two groups come from the same population. To control for the type 1 error rate, the Bonferroni corrected p-values were analyzed. The significance level, alpha, is set to 0.05 for all statistical tests.

## Results

Our primary focus was on the validation of the method by comparing the EEG microstate topographies of patients with disorders of consciousness to those of healthy controls from this study and previous ones. Secondly, we investigate the static microstate markers, specifically the Ratio of Total Time covered (RTT) and Global Explained Variance (GEV). The third part focuses on dynamic microstate metrics, specifically the mean microstate duration (MMD) and microstate duration variance (MDV), transition matrices (TM), and entropy production. The findings highlight microstate differences between healthy controls, patients, and across patient groups.

### EEG Microstates in Patients with Disorders of Consciousness

The EEG microstates depict generalized topographical alterations (Figure 1) whose occurrence and stability are associated with the same or close-by neural sources. The method involves the clustering and segmentation of EEG microstates, where the EEG data is divided into epochs, the Global Field Potential (GFP) peaks are extracted and filtered, and a modified k-means clustering algorithm is applied to identify topographies representing microstate maps, which are then fitted back to the EEG signal to obtain the sequence of dominant maps and their durations. The first question when studying these topographies in patients with disorders of consciousness is whether they will be different from the ones of the healthy controls observed both in this study and previous ones. The final maps include the three-level clustering (Supplementary Figure 3) shown in Figure 1. E, have the same topographies as the ones reported in the literature^6–11,19^. Additionally, the Global Explained Variance (GEV) does not differ between the patient groups and healthy controls (Figure 2. B).

**Figure 2.**
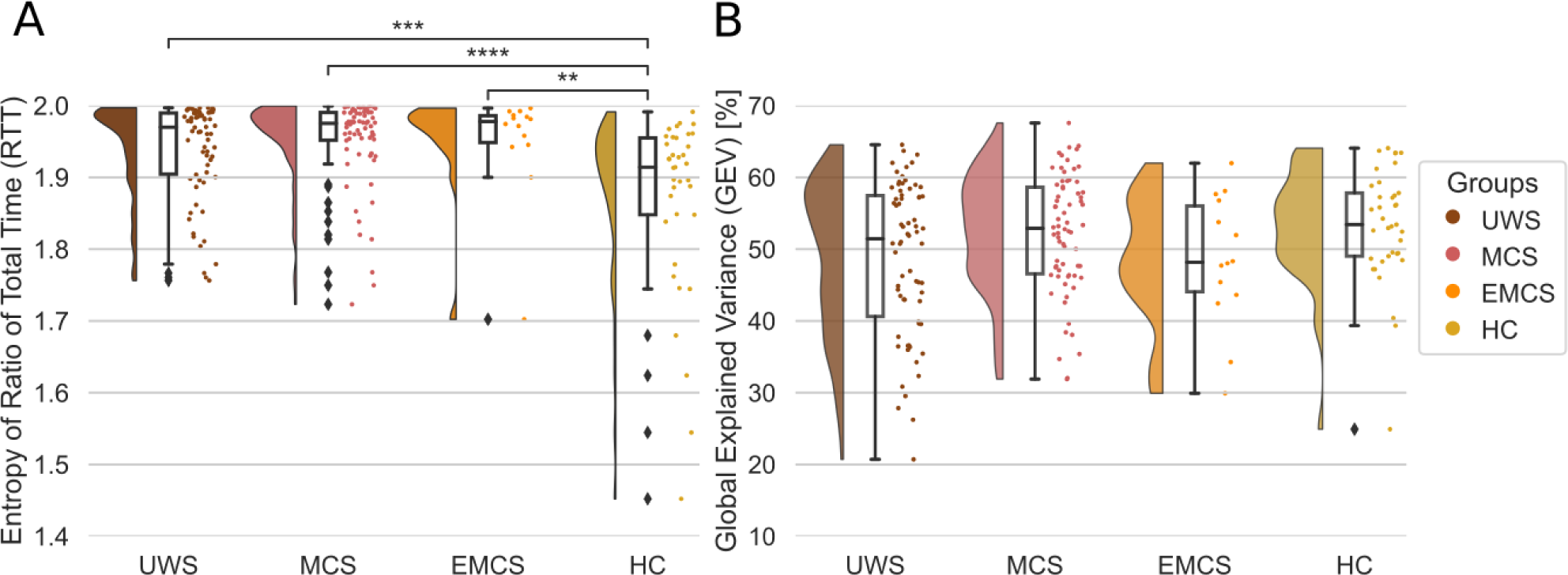
The static microstate metrics Entropy of the Ratio of Total Time (RTT) covered differs between patients with Disorders of Consciousness and Healthy Controls. **(A)** Entropy of the Ratio of Total Time (RTT) covered by the microstates for all three patient groups and the healthy controls. **(B)** Summed Global Explained Variance (GEV) for all microstates together, plotted separately for all three patient groups and the healthy controls. According to the Mann-Whitney U tests, the RTT values between some of the groups are statistically different. All values are Bonferroni corrected. In the figures, one dot represents one subject. The stars represent significance following Mann Whitney U tests between distributions (* p<0.05; ** p<0.01; *** p<0.001; **** p<0.0001). Abbreviations: Ratio of Total Time covered (RTT), Global Explained Variance (GEV), Unresponsive Wakefulness Syndrome (UWS), Minimally Conscious State (MCS), Emergent Minimally Conscious State (EMCS), Healthy Controls (HC).

### Static Microstate Markers

Two measures to assess the static properties of the microstates are the Ratio of Total Time covered (RTT) and Global Explained Variance (GEV) (see Methods). To summarize the occupancy ratios (RTT) of each map, we can calculate their entropy. When looking into the entropy of the Ratio of Total Time covered (RTT) or the ratio of the dominance of one map, in Figure 2 we observe differences between the healthy controls and all the patient groups, but no differences among the patients’ groups (between UWS and HC U(70,37)=1909, p=0.00018; between MCS and HC U(70, 37)=2077, p<0.0001; between EMCS and HC U(14, 37)=425, p=0.0014) The healthy participants, on a group level, show a decrease in entropy, meaning that the occurrences of the four maps are more predictable. Whereas the patient groups have higher entropies showing lower predictability of transitions (or in other words close to uniform probabilities of 25% for the 4 maps). This indicates that some maps, especially in the healthy controls, dominate. To understand this trend we analyzed the RTTs per map and per group (Supplementary Figure 1). We can observe differences in the distributions between HC and the patient groups for maps B and C (map B: MCS & HC U(70,37)=699, p=0.001; map C: UWS & HC U(70,37)=726, p=0.002; MCS & HC U(70,37)=396, p<0.0001; EMCS & HC U(14,37)=82, p=0.002). Map B is less present on average in the HC group, and vice versa for map C. Map D on the other hand shows a bimodal distribution in the HC, where for roughly half of the participants the RTT is as expected (25%) or higher, and for the other half the RTT is much lower. We only see a similar trend between GEV and RTT for map C (Supplementary Figure 1). These differences in RTT in maps B and C for the healthy controls are what contribute to the lower entropy values.

In Figure 2. B the summed GEV of all four maps per participant group is shown. Following Mann-Whitney U two-sided tests, no significant differences between the pairs of groups were observed (p>0.05, Bonferroni corrected). Thus, the ability of the four clustered maps to capture the variance of the true sample-to-sample topographies does not significantly differ between healthy controls and disorders of consciousness patients. In other words, maps do not bias the dynamic markers towards one of the included groups. When we analyze the GEV separately for each map (Supplementary Figure 1. B), we only observe differences between some of the patient groups and the healthy controls in maps B and C (map B: UWS & MCS U(70,70)=1733, p=0.034; MCS & HC U(70,37)=754, p=0.005; map C: UWS & HC U(70,37)=771, p=0.007; MCS & HC U(70,37)=531, p<0.0001; EMCS & HC U(14,37)=104, p=0.013). The inter-group differences reflect the same trends in GEV and RTT which is expected given they revolve around the closeness of the attributed map and the original topography (see Methods). In the literature, similar GEV values are reported^7,8,29^, roughly around 10-15% per microstate map, with higher values, above 20%, for the anterior-posterior microstate C, which is something that we also observe.

### Dynamic Microstate Markers

The dynamic microstate markers show higher inter-group differences (Figure 3). Both the Mean Microstate Durations (MMD) and the Microstate Duration Variances (MDV), on a group level, show a decreasing trend. In other words, going from the patient group UWS to healthy controls, we see a decrease in the microstate durations. Similarly, there is a decrease in the variance of durations which reflects an increased consistency across microstate durations going from UWS to HC. We test this decrease using Mann-Whitney U tests, two-sided and Bonferroni corrected. The differences are significant between most groups for the MMD: UWS and HC U(70,37)=2422, p<0.0001, MCS and HC U(70, 37)=2220, p<0.0001, EMCS and HC U(14, 37)=405, p=0.013; UWS and MCS U(70,70)=3264, p=0.0042; UWS and EMCS U(70,14)=746, p=0.013. For the MDV we observe similar intergroup differences: UWS and HC U(70,37)=2384, p<0.0001, MCS and HC U(70, 37)=2076, p<0.0001; EMCS and HC U(14, 37)=399, p=0.019; UWS and MCS U(70,70)=3303, p=0.002; UWS and EMCS U(70,14)=742, p=0.015. Regarding the MCS and EMCS patient group statistics, a Mann-Whitney U two-sided test shows no significant differences (p>0.05, Bonferroni corrected, U(70,14)=608, p=0.95 for MMD and U(70,14)=565, p=1 for MDV).

**Figure 3.**
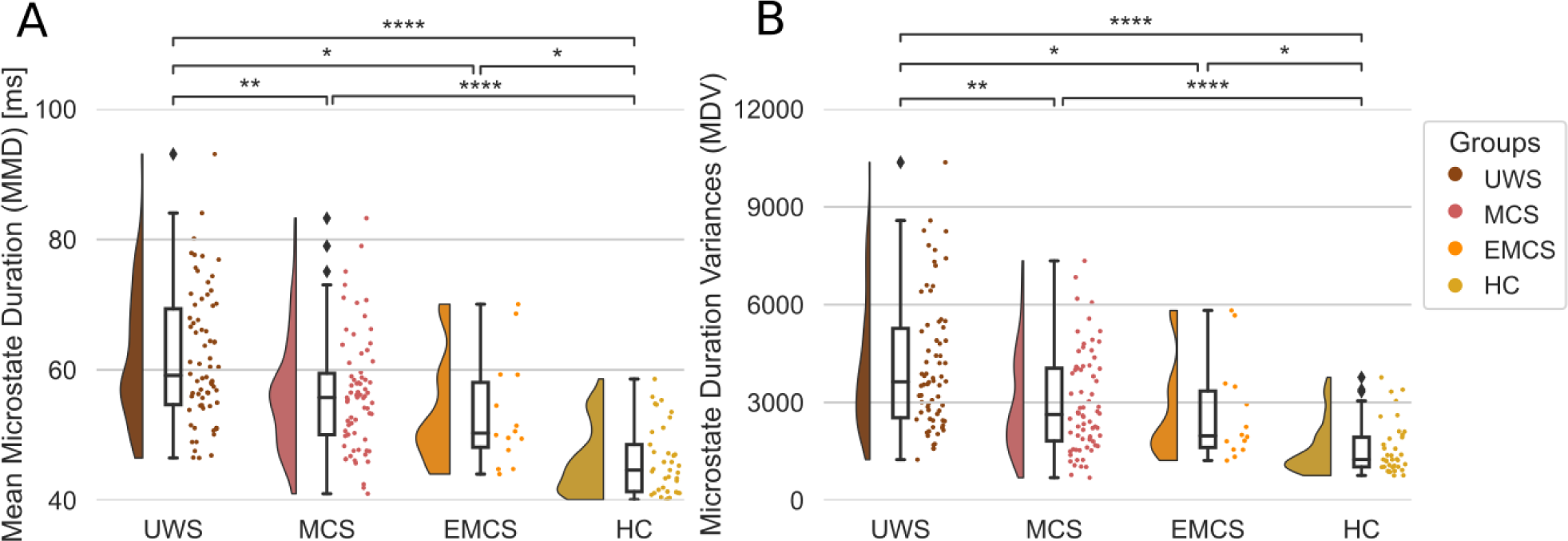
Dynamic microstate markers: MMD and MDV differ between Disorders of Consciousness groups and Healthy Controls. **(A)** Mean Microstate Durations (MMD) for all microstates for all three patient groups and the healthy controls. Statistical differences are found between the healthy controls (HC) and the three patient groups. **(B)** Same as (A) but for the Microstate Duration Variances (MDV). The statistic and the p-value of the Mann-Whitney U two-sided tests for the MMD and the MDV are given in the Results. In all the panels, one dot represents one subject. All p-values are Bonferroni corrected. The stars represent significance following Mann Whitney U tests between distributions (* p<0.05; ** p<0.01; *** p<0.001; **** p<0.0001). Abbreviations: Mean Microstate Duration (MMD), Microstate Duration Variance (MDV), milliseconds (ms), Unresponsive Wakefulness Syndrome (UWS), Minimally Conscious State (MCS), Emergent Minimally Conscious State (EMCS), Healthy Controls (HC).

When we look into the MMD and MDV separately per each map (Supplementary Figure 2), we see that the trend is consistent. For map D, we observe a lower microstate duration especially in the healthy controls as is expected from the lower RTT values of map D for this group. The higher intra-subject variance in the UWS patients reflects instability in the duration of the microstates to a higher degree. Conversely, as HC shows the lowest intra-subject variance, we can state that the microstates’ duration is more consistent in this group and consistently short as shown by the mean microstate durations.

In Figure 4 the average transition probabilities going from map X (row) to map Y (column) are given per group. For the patient groups, we see that more than 90% of the transitions are within the same map, rather than transitioning to another map. This is aligned with the longer durations we observe in the patient groups. In addition, we observe a decrease in symmetry (*P_ij_* ≠ *P_ji_*). This indicates that there are some memory effects in the transitions or non-reversible dynamics^10,11,21^. Another way to represent the information captured by the transition matrices is entropy production^20,21^. The transition matrix entropy production measure per group is given in Figure 4 B where the Mann-Whitney U two-sided tests show significant differences between the healthy controls and the patient groups (UWS and HC U(70,37)=496, p<0.0001; MCS and HC U(70, 37)=447, p<0.0001; EMCS and HC U(14, 37)=90, p=0.001). This shows that the entropy production is higher in healthy subjects than in patients. This indicates that the healthy controls microstate transitions are further from an equilibrium compared to the patient groups.

**Figure 4.**
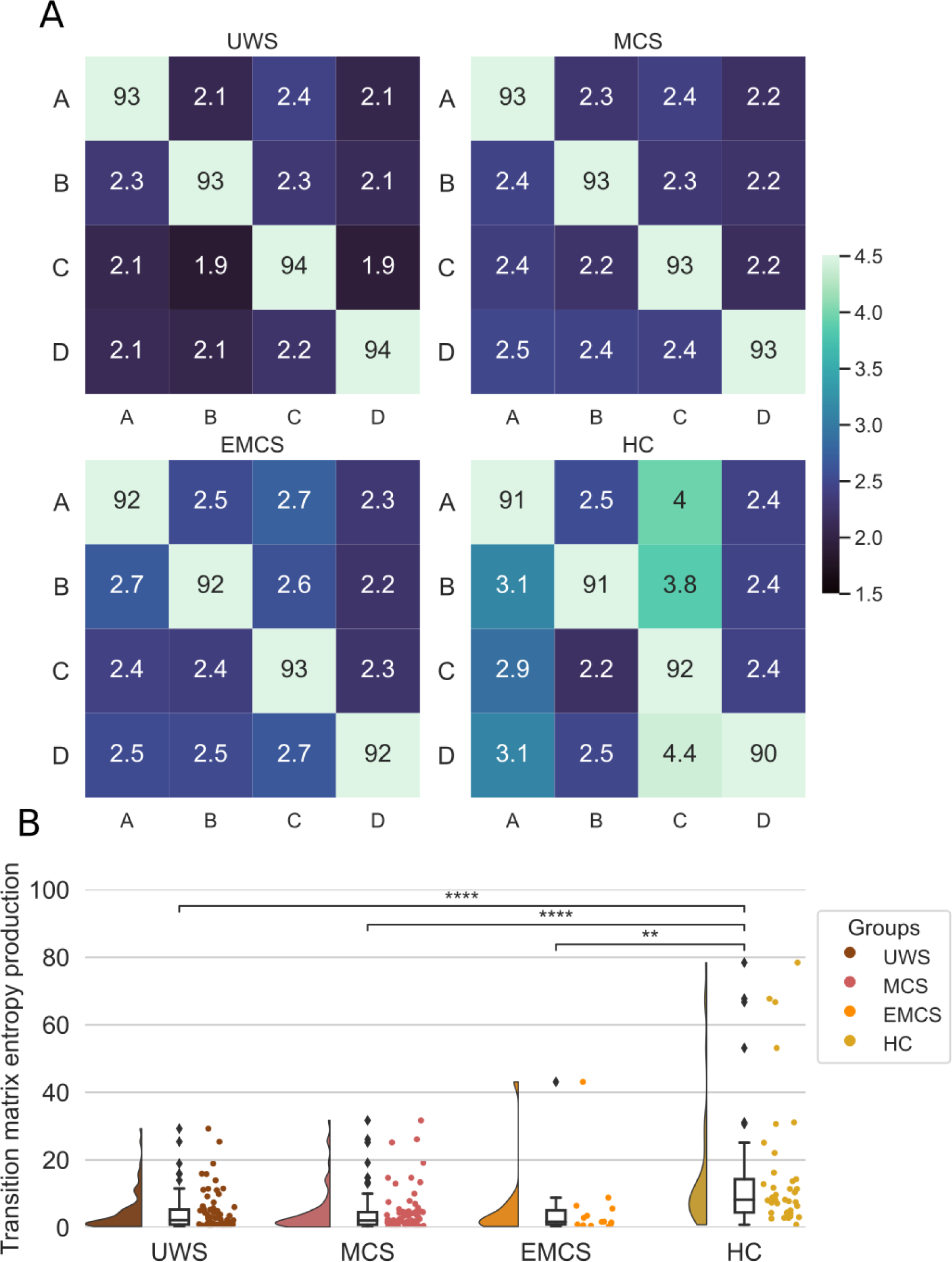
The Microstates Transition Matrices (MTM) summarized using the Entropy productions show differences between Healthy Controls and patients with Disorders of Consciousness. **(A)** The Microstates Transition Matrices (MTM) transitions are shown going from a row value to a column value. The transitions are given in percent. (**B)** Entropy productions per group of the transition probabilities. According to the Mann-Whitney U tests, significant differences were found between the healthy controls and each of the three patient groups. The stars represent significance following Mann Whitney U tests between distributions (* p<0.05; ** p<0.01; *** p<0.001; **** p<0.0001). All values are Bonferroni corrected. In the last panel, one dot represents one subject. Abbreviations: Unresponsive Wakefulness Syndrome (UWS), Minimally Conscious State (MCS), Emergent Minimally Conscious State (EMCS), Healthy Controls (HC).

## Discussion

### Summary

In this study, we used a pseudo-resting-state EEG analysis aimed to investigate the temporal characteristics of ongoing spatial patterns. The configuration of scalp topographies remains semi-stable over successive short-time periods lasting on average from 50 to 100 milliseconds. These periods of stability are what we call microstates Four canonical maps and their temporal properties have been hypothesized to represent the quality of mentation in resting state recordings^6^. However, so far, these topographies have been most often analyzed from low-density EEG recordings. In this study, a high-density (256 electrodes) EEG system was used, which can capture more detailed topographies. In this case, the maps must be precise enough to separate functionally different states (for example in UWS and HC), but also generalizable enough to allow cross-group comparisons^6^. The k-means clustering output resulted in maps that can be reliably compared with the canonical maps typically reported in the literature. A group-level clustering was conducted on all DoC and HC groups together and two groups of markers were investigated, static (RTT and GEV) and dynamic (MMD, MDV, TM, and EP). The markers Ratio of Total Time (RTT) covered, the Global Explained Variance (GEV), and the Mean Microstates Durations (MMD) derived from the HC microstates segmentation, are comparable with other studies^6,8,19,24,35^. We extend the analysis by using metrics that capture different properties of the distributions such as the Microstate Duration Variance (MDV) and Entropy Production (EP).

### Static microstate markers do not differ across patient groups

The static markers, which are not dependent on the temporal dynamics of the microstates, showed no reliable differences among patient group categories (Figure 2). The low GEV of microstate D could reflect a higher heterogeneity of the topographies that were assigned to map D. Such a trend is observed By Brodbeck et al.^7^ but only in non-REM and deep sleep. Interestingly, map D had similar RTT values in UWS and MCS to the ones of other maps, and lower in one part of the HC group. Comsa et al.^8^ findings are aligned with our results, there seems to be a lower entropy, and less balanced RTT when subjects are awake than when they are asleep.

On the contrary, RTT entropy values in our study differ from those observed in another work with patients, where the RTT distributions in HC are more balanced, and less predictable, with a higher entropy; than the UWS and MCS patients^18^. In another work on sleep, similarly, there is a decrease in entropy going from awake to deep sleep, with N2 sleep showing the highest RTT variance between maps (lowest entropy)^7^. However, any comparison between sleep and DoC as states of consciousness has to be taken with precaution because there is mounting evidence that sleep cannot be studied as a uniquely unconscious state, but rather shows high heterogeneity^36^.

### Dynamic microstate markers differ across patient groups

When analyzing the MMD, we observed lower group-level values, going from UWS to MCS, EMCS, and HC. The statistical significance of the Mann-Whitney U two-sided tests confirmed this observation. The group pairwise Mann-Whitney U test, after a Bonferroni correction, revealed significant differences between all groups except for MCS and EMCS (U(70,14)=608, p=0.95). However, due to the unbalanced sample sizes, the power of the test is limited. Furthermore, the shortening of EEG microstates in the HC, reveals faster dynamics in fully preserved consciousness in the healthy controls. This is captured by other studies that investigated EEG microstates in a smaller sample of DoC patients with an average of around 20 ms lengthening of the microstates duration^19^, in drowsy and asleep subjects, with 10 ms MMD lengthening is asleep compared to awake and attentive participants^8^, and 50 ms lengthening in deep sleep compared to wakefulness^7^. In other words, the microstate durations during deep sleep were twice as long compared to when the participants were awake; and the deeper the subjects were asleep, the slower the topographical alterations were. Furthermore, the average duration of microstate D was significantly increased when participants in the study were drowsy or had fallen asleep in comparison to when they were awake^8^. A parallel can be drawn between the studies done on sleep and DoC patients. Sleep, which is a transient loss of consciousness, and DoC classes which reflect a non-reversible loss of consciousness condition, both show similar microstate dynamics. This implies that the microstate temporal dynamics correlate with wakefulness or lack thereof.

Similarly, in the Microstate Duration Variance (MDV) distributions, a lowering of per-subject variance scores from UWS to HC was observed. The group pairwise Mann-Whitney U tests, after a Bonferroni correction, revealed significant differences between all pairs of groups except for MCS and EMCS (U(70,14)=565, p=1) (Figure 3). The initial hypothesis we postulated is that the HC group will show the highest per-subject variance. The line of thought is that the consciousness level, and thus inter-subject differences would be reflected in a higher variance of microstate durations. A variance that should not show up in patient groups. Similar variances (spread) of the MMD distributions are reported when participants are asleep compared to when they are awake^7,8^, especially when comparing deep (N3) sleep compared to wakefulness^7^. However, in both studies, the subject-level MDV is not calculated, and a direct comparison entails further investigation.

When looking into the lower per-subject variance in healthy controls, one could postulate that the microstates do not reflect inter-subject conscious content, but rather more general brain dynamics of conscious states. Furthermore, the higher variance in the patient groups could reflect the high heterogeneity of the etiologies and clinical pictures across patients. Another possibility is that in the patient groups, the microstate durations are consistently long, but because we use short epochs, their durations are interrupted.

### Entropy production is lower in patients compared to healthy controls

The decrease in symmetrical transitions across states is captured by the measure of entropy production^20,21^. On a group level, the differences are significant between the healthy controls and the three patient groups. This indicates that the microstates time series in the healthy controls has higher irreversibility, or there is a breaking in detailed balance. In contrast, the patient groups show entropy production values aligned with higher time reversibility. In each of the groups, there are a few patients who show higher state transition asymmetry. Further investigations can test if the higher entropy production in patients is aligned with a better prognostic^20,21^. Our results are aligned with previous observations which report higher entropy production with an increase in task cognitive demand^21^ and a decrease in global states of unconsciousness^20^. Other analogous metrics have been proposed, where a breaking of temporal symmetry (analogous to the entropy production) is shown in healthy control compared to patients with a disorder of consciousness^37^.

### Conclusions

In this study, we explored the dynamics of EEG microstates in patients with disorders of consciousness (DoC) to understand residual brain activity and the reorganization of brain networks on a sub-second scale. By analyzing static and dynamic EEG microstate markers, we aimed to differentiate between healthy controls and DoC patients and among different DoC groups. Our findings indicate that while static markers like the Ratio of Total Time covered (RTT) and Global Explained Variance (GEV) do not distinguish between patient groups, dynamic markers reveal significant inter-group differences. Specifically, the Mean Microstate Durations (MMD) and Microstate Duration Variances (MDV) decrease with higher consciousness levels, whereas non-diagonal transitions in Microstate Transition Matrices (MTM) and Entropy Production (EP) increase. These results suggest that DoC patients exhibit slower and more equilibrium-like brain dynamics, reflecting a state closer to time-reversibility. This study enhances our understanding of brain dynamics in DoC patients by showing that dynamic EEG microstate metrics are more sensitive in capturing subtle differences in brain activity among patients with varying levels of consciousness, and translates entropy production measures to EEG acquisitions. Further lines of research can investigate the dynamic EEG microstate markers in movie watching or active tasks, where the differences across patient groups could be further enhanced, as expected from fMRI studies in healthy controls^21^.

## Abbreviations

(DoC): Disorders of Consciousness
(EEG): Electroencephalography
(RTT): Ratio of Total Time covered
(MMD): Mean Microstate Durations
(MDV): Microstate Duration Variances
(MTM): Microstate Transition Matrices
(EP): Entropy Production
(UWS): Unresponsive Wakefulness Syndrome
(VS): Vegetative State
(MCS): Minimally Conscious State
(EMCS): Emergent Minimally Conscious State
(GFP): Global Field Power
(GEV): Global Explained Variance
(GMD): Global Map Dissimilarity
(HC): Healthy Controls
(fMRI): Functional Magnetic Resonance Imaging
(TM): Transition Matrices
(LG) Paradigm: Local-Global

## Data Availability

The data is not publicly available.

For the analyses, we used open-source software MNE Python and the associated implementation of the EEG microstates analysis (https://github.com/wmvanvliet/mne_microstates).

## Acknowledgments

We thank all of the participants who took part in the studies. We would like to thank the work and support of the clinicians at the Neuro ICU, DMU Neurosciences, APHP- Sorbonne Université, Hôpital de la Pitié Salpêtrière, Paris, France; and the patient families whose consent and understanding are essential to the progress of the field.

## Supplementary Materials

**Supplementary Figure 1.**
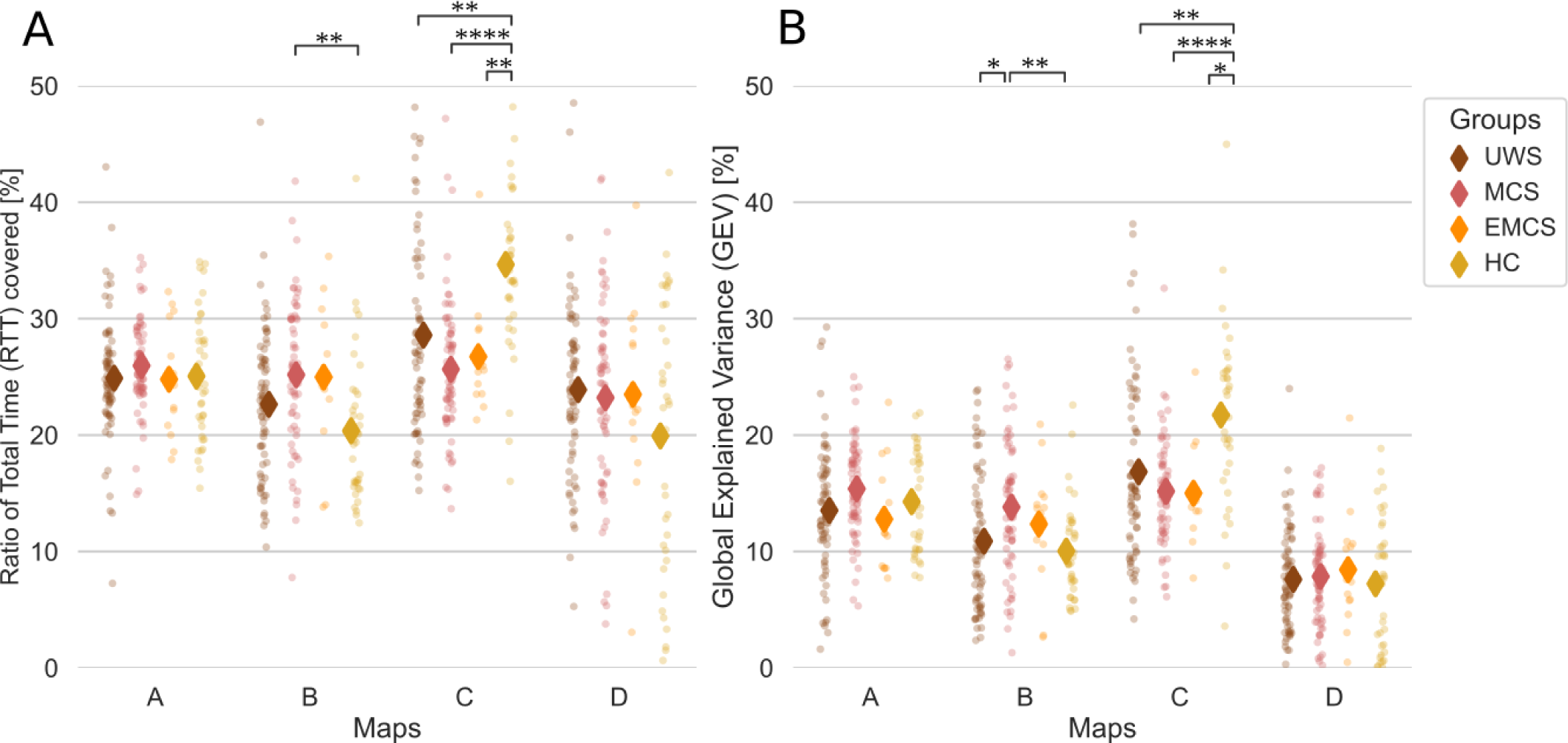
Per-map distributions of the Ratio of Total Time covered (RTT) and Global Explained Variance (GEV) **(A)** The RTT covered are shown separately for all of the different microstates (A, B, C, D). The RTT reflects the percentage of time the given microstate is dominant over the whole duration of the data. **(B)** Same as (A) but for the GEV. The statistic and the p-value of the Mann-Whitney U two-sided tests for the RTT and the GEV per map are given in Supplementary Table 1. All values are Bonferroni corrected. One dot represents one subject. The stars represent significance following Mann Whitney U tests between distributions (* p<0.05; ** p<0.01; *** p<0.001; **** p<0.0001). Abbreviations: Ratio of Total Time covered (RTT), Global Explained Variance (GEV), Unresponsive Wakefulness Syndrome (UWS), Minimally Conscious State (MCS), Emergent Minimally Conscious State (EMCS), Healthy Controls (HC).

**Supplementary Table 1.** Statistics and the p-value of the Mann-Whitney U two-sided tests, Bonferroni corrected, for the Ratio of Total Time covered (RTT) and the Global Explained Variance (GEV) per group per map. The values in bold pass the significance threshold of p<0.05. Abbreviations: Ratio of Total Time covered (RTT), Global Explained Variance (GEV), Unresponsive Wakefulness Syndrome (UWS), Minimally Conscious State (MCS), Emergent Minimally Conscious State (EMCS), Healthy Controls (HC).

| Map | Group 1 | Group 2 | RTT<br>(statistic, p-value) | GEV<br>(statistic, p-value) |
| --- | --- | --- | --- | --- |
| A | UWS | MCS | U(70,70)=1992, $p=0.679$ | U(70,70)=1833, $p=0.122$ |
| A | UWS | EMCS | U(70,14)=468, $p=1$ | U(70,14)=423, $p=1$ |
| A | UWS | HC | U(70,37)=1278, $p=1$ | U(70,37)=1175, $p=1$ |
| A | MCS | EMCS | U(70,14)=409, $p=1$ | U(70,14)=307, $p=0.342$ |
| A | MCS | HC | U(70,37)=1124, $p=1$ | U(70,37)=1109, $p=1$ |
| A | EMCS | HC | U(14,37)=257, $p=1$ | U(14,37)=201, $p=1$ |
| B | UWS | MCS | U(70,70)=1830, $p=0.118$ | **U(70,70)=1733, $p=0.034$** |
| B | UWS | EMCS | U(70,14)=371, $p=1$ | U(70,14)=392, $p=1$ |
| B | UWS | HC | U(70,37)=982, $p=0.488$ | U(70,37)=1248, $p=1$ |
| B | MCS | EMCS | U(70,14)=471, $p=1$ | U(70,14)=428, $p=1$ |
| B | MCS | HC | **U(70,37)=699, $p=0.001$** | **U(70,37)=754, $p=0.005$** |
| B | EMCS | HC | U(14,37)=139, $p=0.14$ | U(14,37)=165, $p=0.581$ |
| C | UWS | MCS | U(70,70)=2059, $p=1$ | U(70,70)=2273, $p=1$ |
| C | UWS | EMCS | U(70,14)=456, $p=1$ | U(70,14)=437, $p=1$ |
| C | UWS | HC | **U(70,37)=726, $p=0.002$** | **U(70,37)=771, $p=0.007$** |
| C | MCS | EMCS | U(70,14)=426, $p=1$ | U(70,14)=459, $p=1$ |
| C | MCS | HC | **U(70,37)=396, $p<0.0001$** | **U(70,37)=531, $p<0.0001$** |
| C | EMCS | HC | **U(14,37)=82, $p=0.002$** | **U(14,37)=104, $p=0.013$** |
| D | UWS | MCS | U(70,70)=2366, $p=1$ | U(70,70)=2311, $p=1$ |
| D | UWS | EMCS | U(70,14)=490, $p=1$ | U(70,14)=448, $p=1$ |
| D | UWS | HC | U(70,37)=1096, $p=1$ | U(70,37)=1227, $p=1$ |
| D | MCS | EMCS | U(70,14)=470, $p=1$ | U(70,14)=461, $p=1$ |
| D | MCS | HC | U(70,37)=1105, $p=1$ | U(70,37)=1193, $p=1$ |
| D | EMCS | HC | U(14,37)=232, $p=1$ | U(14,37)=229, $p=1$ |

**Supplementary Figure 2.**
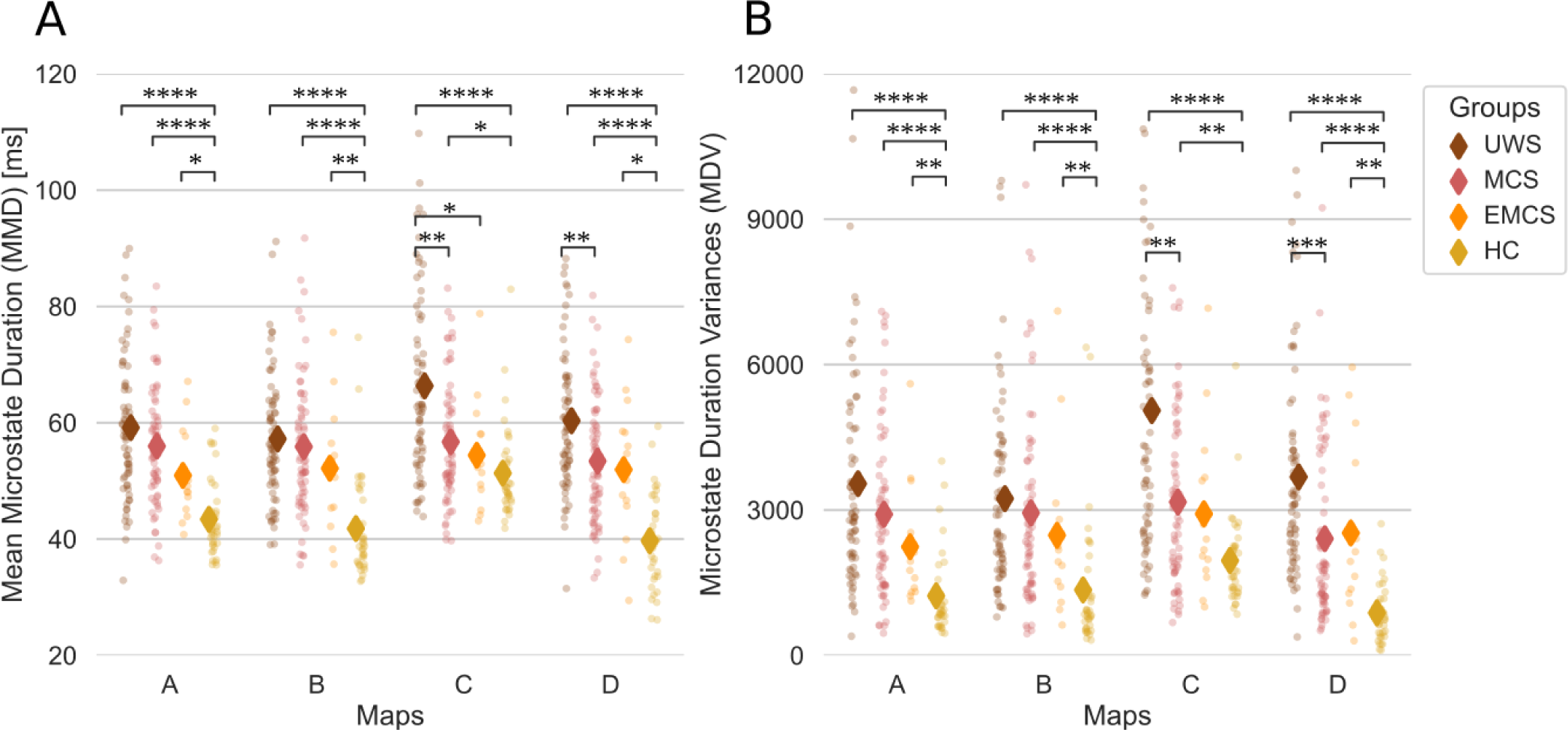
Per-map distributions of the Mean Microstate Durations (MMD) and the Microstate Duration Variances (MDV) **(A)** The MMD are shown separately for all of the different microstates (A, B, C, D). **(B)** Same as (A) but for the MDV. The statistic and the p-value of the Mann-Whitney U two-sided tests for the MMD and the MDV per map are given in Supplementary Table 2. All values are Bonferroni corrected. One dot represents one subject. The stars represent significance following Mann Whitney U tests between distributions (* p<0.05; ** p<0.01; *** p<0.001; **** p<0.0001). Abbreviations: Mean Microstate Duration (MMD), Microstate Duration Variance (MDV), milliseconds (ms), Unresponsive Wakefulness Syndrome (UWS), Minimally Conscious State (MCS), Emergent Minimally Conscious State (EMCS), Healthy Controls (HC).

**Supplementary Table II.** Statistics and the p-value of the Mann-Whitney U two-sided tests, Bonferroni corrected, for the Mean Microstate Durations (MMD) and the Microstate Duration Variances (MDV) per group per map. The values in bold pass the significance threshold of p<0.05. Abbreviations: Mean Microstate Duration (MMD), Microstate Duration Variance (MDV), milliseconds (ms), Unresponsive Wakefulness Syndrome (UWS), Minimally Conscious State (MCS), Emergent Minimally Conscious State (EMCS), Healthy Controls (HC)

| Map | Group 1 | Group 2 | MMD<br>(statistic, p-value) | MDV<br>(statistic, p-value) |
| --- | --- | --- | --- | --- |
| A | UWS | MCS | U(70,70)=2092, $p=1$ | U(70,70)=2059, $p=1$ |
| A | UWS | EMCS | U(70,14)=274, $p=0.116$ | U(70,14)=288, $p=0.187$ |
| A | UWS | HC | **U(70,37)=276, $p < 0.0001$** | **U(70,37)=280, $p < 0.0001$** |
| A | MCS | EMCS | U(70,14)=328, $p=0.631$ | U(70,14)=363, $p=1$ |
| A | MCS | HC | **U(70,37)=365, $p < 0.0001$** | **U(70,37)=409, $p < 0.0001$** |
| A | EMCS | HC | **U(14,37)=102, $p=0.012$** | **U(14,37)=81, $p=0.002$** |
| B | UWS | MCS | U(70,70)=2253, $p=1$ | U(70,70)=2182, $p=1$ |
| B | UWS | EMCS | U(70,14)=359, $p=1$ | U(70,14)=357, $p=1$ |
| B | UWS | HC | **U(70,37)=270, $p < 0.0001$** | **U(70,37)=365, $p < 0.0001$** |
| B | MCS | EMCS | U(70,14)=399, $p=1$ | U(70,14)=411, $p=1$ |
| B | MCS | HC | **U(70,37)=358, $p < 0.0001$** | **U(70,37)=470, $p < 0.0001$** |
| B | EMCS | HC | **U(14,37)=98, $p=0.009$** | **U(14,37)=112, $p=0.024$** |
| C | UWS | MCS | **U(70,70)=1550, $p=0.002$** | **U(70,70)=1511, $p=0.001$** |
| C | UWS | EMCS | **U(70,14)=248, $p=0.045$** | U(70,14)=261, $p=0.073$ |
| C | UWS | HC | **U(70,37)=455, $p < 0.0001$** | **U(70,37)=331, $p < 0.0001$** |
| C | MCS | EMCS | U(70,14)=426, $p=1$ | U(70,14)=449, $p=1$ |
| C | MCS | HC | **U(70,37)=823, $p=0.024$** | **U(70,37)=738, $p=0.003$** |
| C | EMCS | HC | U(14,37)=201, $p=1$ | U(14,37)=168, $p=0.673$ |
| D | UWS | MCS | **U(70,70)=1608, $p=0.005$** | **U(70,70)=1427, $p=0.0002$** |
| D | UWS | EMCS | U(70,14)=314, $p=0.422$ | U(70,14)=312, $p=0.398$ |
| D | UWS | HC | **U(70,37)=156, $p < 0.0001$** | **U(70,37)=110, $p < 0.0001$** |
| D | MCS | EMCS | U(70,14)=460, $p=1$ | U(70,14)=454, $p=0.335$ |
| D | MCS | HC | **U(70,37)=380, $p < 0.0001$** | **U(70,37)=409, $p < 0.0001$** |
| D | EMCS | HC | **U(14,37)=107, $p=0.017$** | **U(14,37)=90, $p=0.005$** |

**Supplementary Figure 3.**
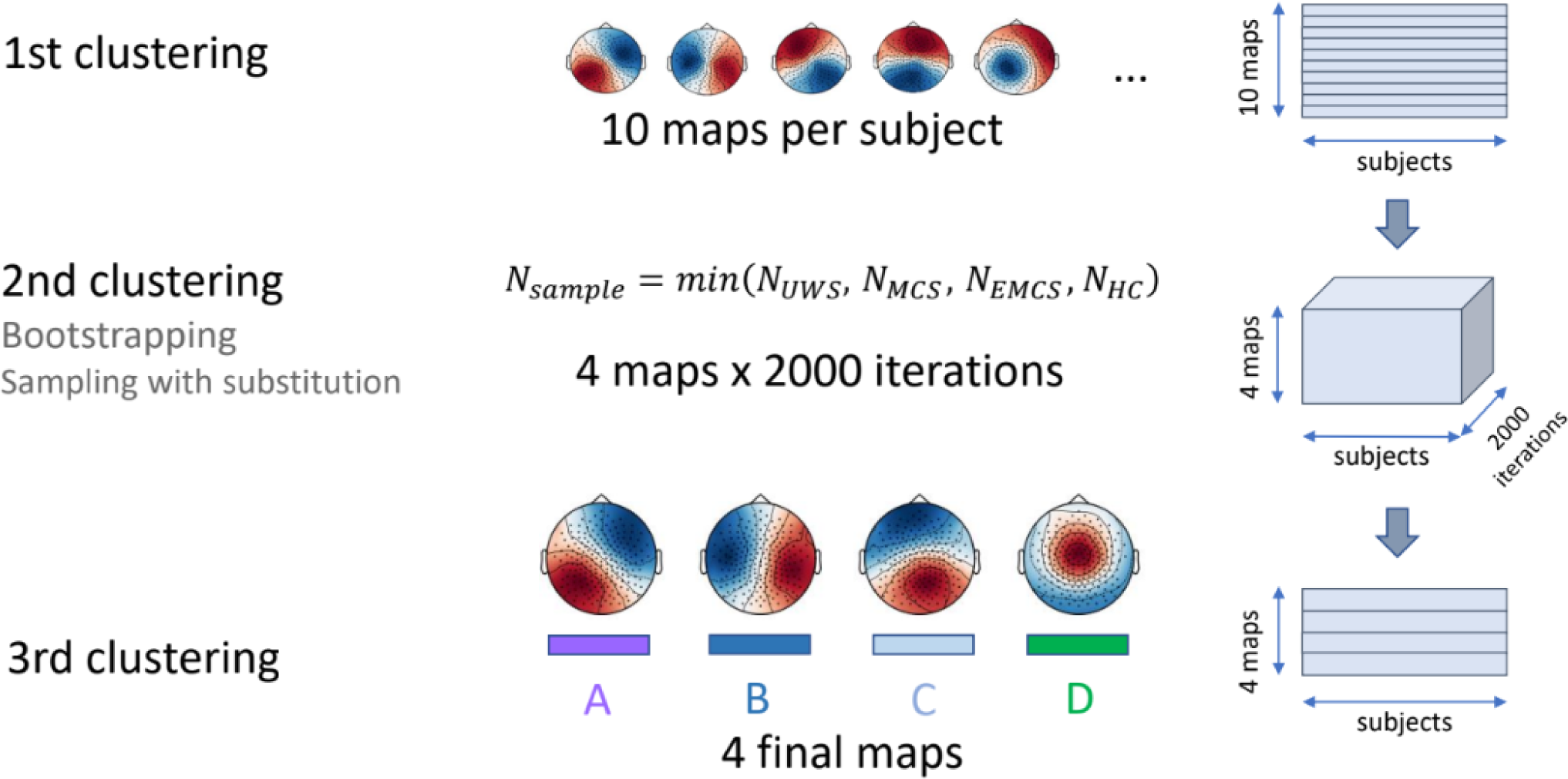
Graphical representation of the 3-level clustering. In the first level, we set the number of clusters k to 10. This is done on a single subject level. The 10 resulting maps from the modified k-means clustering are fed to a second-level clustering where k=4. In this step, we do a bootstrapping due to the unbalanced sample sizes. We take samples with repetition from each group with a sample number equal to the largest sample size per group - in our case N_EMCS_=14. We do this clustering on the sub-samples 2000 times. Thus we obtain 4 x 2000 maps, which we give to a 3rd-level clustering where k=4. These last 4 maps are back-fitted to the time series and thus the segmentation of the microstates is obtained. Abbreviations: Unresponsive Wakefulness Syndrome (UWS), Minimally Conscious State (MCS), Emergent Minimally Conscious State (EMCS), Healthy Controls (HC).

## References

1. Seth AK, Bayne T. Theories of consciousness. Nat Rev Neurosci. 2022;23(7):439-452. doi:10.1038/s41583-022-00587-4

2. Bodien YG, Chatelle C, Edlow BL. Functional Networks in Disorders of Consciousness. Semin Neurol. 2017;37(5):485–502. doi:10.1055/S-0037-1607310

3. Demertzi A, Tagliazucchi E, Dehaene S, et al. Human consciousness is supported by dynamic complex patterns of brain signal coordination. Sci Adv. 2019;5(2):eaat7603. doi:10.1126/sciadv.aat7603

4. Deco G, Jirsa VK, McIntosh AR. Emerging concepts for the dynamical organization of resting-state activity in the brain. Nat Rev Neurosci. 2011;12(1):43–56. doi:10.1038/nrn2961

5. Lehmann D. Principles of spatial analysis. Handb Electroencephalogr Clin Neurophysiol Methods Anal Brain Electr Magn Signals. 1987;1:309–354.

6. Michel CM, Koenig T. EEG microstates as a tool for studying the temporal dynamics of whole-brain neuronal networks: A review. NeuroImage. 2018;180:577–593. doi:10.1016/j.neuroimage.2017.11.062

7. Brodbeck V, Kuhn A, von Wegner F, et al. EEG microstates of wakefulness and NREM sleep. NeuroImage. 2012;62(3):2129–2139. doi:10.1016/j.neuroimage.2012.05.060

8. Comsa IM, Bekinschtein TA, Chennu S. Transient Topographical Dynamics of the Electroencephalogram Predict Brain Connectivity and Behavioural Responsiveness During Drowsiness. Brain Topogr. 2019;32(2):315–331. doi:10.1007/s10548-018-0689-9

9. Kuhn A, Brodbeck V, Tagliazucchi E, Morzelewski A, von Wegner F, Laufs H. Narcoleptic Patients Show Fragmented EEG-Microstructure During Early NREM Sleep. Brain Topogr. 2015;28(4):619–635. doi:10.1007/s10548-014-0387-1

10. von Wegner F, Tagliazucchi E, Laufs H. Information-theoretical analysis of resting state EEG microstate sequences - non-Markovianity, non-stationarity and periodicities. NeuroImage. 2017;158:99–111. doi:10.1016/j.neuroimage.2017.06.062

11. von Wegner F, Laufs H. Information-Theoretical Analysis of EEG Microstate Sequences in Python. Front Neuroinformatics. 2018;12:30. doi:10.3389/fninf.2018.00030

12. Stefan S, Schorr B, Lopez-Rolon A, et al. Consciousness Indexing and Outcome Prediction with Resting-State EEG in Severe Disorders of Consciousness. Brain Topogr. 2018;31(5):848–862. doi:10.1007/s10548-018-0643-x

13. Britz J, Van De Ville D, Michel CM. BOLD correlates of EEG topography reveal rapid resting-state network dynamics. NeuroImage. 2010;52(4):1162–1170. doi:10.1016/j.neuroimage.2010.02.052

14. Bekinschtein TA, Dehaene S, Rohaut B, Tadel F, Cohen L, Naccache L. Neural signature of the conscious processing of auditory regularities. Proc Natl Acad Sci. 2009;106(5):1672–1677. doi:10.1073/pnas.0809667106

15. Jennett B, Plum F. Persistent Vegetative State after brain damage: A Syndrome in Search of a Name. The Lancet. 1972;299(7753):734-737. doi:10.1016/S0140-6736(72)90242-5

16. Giacino JT, Ashwal S, Childs N, et al. The minimally conscious state: definition and diagnostic criteria. Neurology. 2002;58(3):349–353. doi:10.1212/WNL.58.3.349

17. Giacino JT, Schnakers C, Rodriguez-Moreno D, Kalmar K, Schiff N, Hirsch J. Behavioral assessment in patients with disorders of consciousness: gold standard or fool’s gold? Prog Brain Res. 2009;177:33–48. doi:10.1016/S0079-6123(09)17704-X

18. Gui P, Jiang Y, Zang D, et al. Assessing the depth of language processing in patients with disorders of consciousness. Nat Neurosci. 2020;23(6):761–770. doi:10.1038/s41593-020-0639-1

19. Toplutaş E, Aydın F, Hanoğlu L. EEG Microstate Analysis in Patients with Disorders of Consciousness and Its Clinical Significance. Brain Topogr. 2024;37(3):377–387. doi:10.1007/S10548-023-00939-Y/FIGURES/5

20. Sanz Perl Y, Bocaccio H, Pallavicini C, et al. Nonequilibrium brain dynamics as a signature of consciousness. Phys Rev E. 2021;104(1). doi:10.1103/PhysRevE.104.014411

21. Lynn CW, Cornblath EJ, Papadopoulos L, Bertolero MA, Bassett DS. Broken detailed balance and entropy production in the human brain. PNAS. Published online May 5, 2021. doi:10.1073/pnas.2109889118

22. Engemann DA, Raimondo F, King JR, et al. Robust EEG-based cross-site and cross-protocol classification of states of consciousness. Brain. 2018;141(11):3179–3192. doi:10.1093/brain/awy251

23. Sitt JD, King JR, El Karoui I, et al. Large scale screening of neural signatures of consciousness in patients in a vegetative or minimally conscious state. Brain. 2014;137(8):2258–2270. doi:10.1093/brain/awu141

24. Gramfort A, Luessi M, Larson E, et al. MNE software for processing MEG and EEG data. NeuroImage. 2014;86:446–460. doi:10.1016/j.neuroimage.2013.10.027

25. Milz P, Faber PL, Lehmann D, Koenig T, Kochi K, Pascual-Marqui RD. The functional significance of EEG microstates—Associations with modalities of thinking. NeuroImage. 2016;125:643–656. doi:10.1016/j.neuroimage.2015.08.023

26. Sikka A, Jamalabadi H, Krylova M, et al. Investigating the temporal dynamics of electroencephalogram (EEG) microstates using recurrent neural networks. Hum Brain Mapp. Published online February 24, 2020:hbm.24949. doi:10.1002/hbm.24949

27. Michel CM, Koenig T, Brandeis D, Gianotti LRR, Wackermann J. Electrical Neuroimaging. Cambridge Medicine; 2009.

28. Poulsen AT, Pedroni A, Langer N, Hansen LK. Microstate EEGlab toolbox: An introductory guide. bioRxiv. Published online 2018:1–30. doi:10.1101/289850

29. Zanesco AP, King BG, Skwara AC, Saron CD. Within and between-person correlates of the temporal dynamics of resting EEG microstates. NeuroImage. 2020;211:116631. doi:10.1016/j.neuroimage.2020.116631

30. Pascual-Marqui RD, Michel CM, Lehmann D. Segmentation of brain electrical activity into microstates: model estimation and validation. IEEE Trans Biomed Eng. 1995;42(7):658–665. doi:10.1109/10.391164

31. Mishra A, Englitz B, Cohen MX. EEG microstates as a continuous phenomenon. NeuroImage. 2020;208:116454. doi:10.1016/j.neuroimage.2019.116454

32. Shaw SB, Dhindsa K, Reilly JP, Becker S. Capturing the Forest but Missing the Trees: Microstates Inadequate for Characterizing Shorter-Scale EEG Dynamics. Neural Comput. 2019;31(11):2177–2211. doi:10.1162/neco_a_01229

33. Murray MM, Brunet D, Michel CM. Topographic ERP Analyses: A Step-by-Step Tutorial Review. Brain Topogr. 2008;20(4):249–264. doi:10.1007/s10548-008-0054-5

34. Titterington DM, Titterington P of SDM, Smith AFM, Makov UE. Statistical Analysis of Finite Mixture Distributions. Wiley; 1985.

35. Koenig T, Prichep L, Lehmann D, et al. Millisecond by Millisecond, Year by Year: Normative EEG Microstates and Developmental Stages. NeuroImage. 2002;16(1):41–48. doi:10.1006/nimg.2002.1070

36. Andrillon T, Poulsen AT, Hansen LK, L É Ger D, Kouider S. Neural markers of responsiveness to the environment in human sleep. J Neurosci. 2016;36(24):6583–6596. doi:10.1523/JNEUROSCI.0902-16.2016

37. Guzmán EG, Perl YS, Vohryzek J, et al. The lack of temporal brain dynamics asymmetry as a signature of impaired consciousness states. Interface Focus. 2023;13(3). doi:10.1098/RSFS.2022.0086

